# Male-Specific Protein Disulphide Isomerase Function is Essential for *Plasmodium* Fertilization and Transmission

**DOI:** 10.1101/411926

**Authors:** Fiona Angrisano, Katarzyna A. Sala, Sofia Tapanelli, George K. Christophides, Andrew M. Blagborough

## Abstract

Inhibiting transmission of *Plasmodium* is an essential strategy in malaria eradication, and the biological process of gamete fusion during fertilization is a proven target for this approach. The lack of knowledge of the mechanisms underlying fertilization have been a hindrance in the development of transmission-blocking interventions. Here we describe a protein disulphide isomerase essential for malarial transmission (*PDI-Trans*/PBANKA_0820300) to the mosquito. We show that *PDI-Trans* activity is male-specific, surface expressed, essential for fertilization/transmission, and exhibits disulphide isomerase activity which is up-regulated post-gamete activation. We demonstrate that *PDI-Trans* is a viable anti-malarial drug and vaccine target blocking malarial transmission with the use of the PDI inhibitor bacitracin (98.21%/92.48% reduction in intensity/prevalence), and anti-*PDI-Trans* peptide antibodies (66.22%/33.16% reduction in intensity/prevalence). To our knowledge, these results provide the first primary evidence that protein disulphide isomerase function is essential for malarial transmission, and emphasize the potential of anti-PDI agents to act as anti-malarials, facilitating the future development of novel transmission-blocking compounds or vaccines.

## Introduction

Malaria remains a major global health challenge with an estimated 216 million new cases and 445,000 deaths in 2016 [1]. Current tools have substantially reduced the global burden of disease, but recent progress has stalled [1], and it is widely accepted that a range of new tools will be needed to achieve malaria elimination [2]. The causative agent of malaria, the protozoan parasite *Plasmodium*, is transmitted almost exclusively by mosquitoes of the genus *Anopheles*. Transmission of *Plasmodium* from humans to mosquitoes is entirely dependent on the presence of sexually committed gametocytes in circulating blood, which rapidly undergo the process of activation and differentiate into male (micro) and female (macro) gametes upon uptake by the mosquito within a blood meal. The essential process of fertilization is then initiated by gamete adhesion, followed by membrane fusion [3,4]. A small number of proteins have been previously been implicated in plasmodial fertilization; the 6-Cys protein family members P48/45, P47 and P230 have been shown to have a demonstrable role in the mutual recognition and adhesion of micro- and macro-gametes [5-7], whereas the conserved male-specific Class II fusion protein HAP2/GCS1 has been shown to be the key driver of membrane fusion by mediating merger of lipid bilayers [3-4]. Following successful fertilization, resulting zygotes develop into motile ookinetes, establishing infection in the insect host by migration and invasion of the mosquito midgut, allowing for the progression of the parasitic lifecycle. Despite the obvious biological importance of parasitic transmission and its proven previous targeting as a potential point to disrupt the parasitic lifecycle with multiple therapeutics [8], our knowledge of the cellular and molecular mechanisms underlying fertilization and subsequent zygote formation in *Plasmodium* are surprisingly sparse.

It is widely recognized that to achieve malarial eradication, it will be necessary to use interventions that inhibit the transmission of parasites from humans to mosquitoes [2]. A potential manner of achieving this is by targeting *Plasmodium* using transmission-blocking interventions (TBIs); i.e. transmission blocking vaccines (TBVs), or transmission blocking drugs (TBDs) against parasitic sexual stages [9]. Antibodies targeted to three of the five currently proven, potent TBV targets have confirmed localization to proteins found on the surface of the plasma membrane of the gametes [10-20], clearly indicating the potential value of targeting this stage of the parasite lifecycle. Additionally, multiple anti-malarial compounds have been demonstrated to have activity against this parasitic stage [21-25]. In summary, the comparatively short life span, increased fragility and availability of proven surface-localized proteins on the male gamete of *Plasmodium* make targeting this gamete stage of the lifecycle a potential method of inhibiting transmission [26,27]. Similarly, potent TBIs targeting the parasitic ookinete post-fertilization are well characterized in multiple vaccine and drug studies [16,17,25, 28-30].

Protein Disulphide Isomerase (PDI) (EC: 5.3.4.1) is a multifunctional member of the thioredoxin superfamily of redox proteins, characterized by the presence of the βαβαβαββα fold [31]. PDIs typically have three catalytic activities; disulphide isomerase, thiol-disuphide oxidoredctase, and redox-dependent chaperone. PDI homologues have been identified in multiple species, where they are “classically” located in the endoplasmic reticulum (ER) and facilitate the folding and assembly of secretory and membrane proteins within the lumen [32]. In *Plasmodium*, a small number of proteins have been putatively identified (by sequence homology) as PDI-like molecules in *Plasmodium falciparum, vivax, knowlesi, berghei* and *yoelii* [33,34]. Conclusive demonstration of PDI activity has currently only been demonstrated with PF3D7_0827900/PDI-8 [33], with transcription and translation demonstrated in asexual blood schizonts, gametocytes and sporozoites. Knowledge regarding the process of disulphide bond-dependent protein folding in *Plasmodium* is scarce. Similarly, to target the sexual stages of the malaria parasite further, a deeper understanding of transmission and specifically, the mechanism of fertilization within *Plasmodium* is vital, and offers the potential for the development of new, effective anti-malarial TBIs. Here, we describe the identification, characterization and role of a protein disulphide isomerase essential for malarial transmission (*PDI-Trans*/PBANKA_0820300) to the mosquito host in *P. berghei*. We demonstrate that *PDI-Trans* is transcribed and translated across the entire parasitic lifecycle, but exhibits activity at the sexual stages of the lifecycle, when fertilization of gametes occurs. We show that *PDI-Trans* is male specific, essential for successful fertilization/transmission, and exhibits disulphide isomerase function which is up-regulated post-gamete activation. Furthermore, we show that *PDI-Trans* is a viable anti-malarial drug and vaccine target, expressed on the surface of the sexual stages of *Plasmodium*, by blocking malarial transmission with the use of repurposed compounds that target PDI activity, and anti-*PDI-Trans* peptide antibodies. These results demonstrate that protein disulphide isomerase function is essential for malarial transmission, emphasize the potential of anti-PDI agents to act as anti-malarials, and demonstrate the potential utility of rationally selected targets to facilitate the development of novel anti-malarial transmission-blocking interventions.

## Results

### PDI-Trans is located on the surface on the transmission stages of P. berghei

Previous proteomic analysis of a *P. berghei* male gamete proteome generated in [35] followed by advanced bioinformatics analysis encompassing a suite of functional and localization-based algorithms [36] identified the expression of *PDI-Trans* (PBANKA_0820300) in the male gamete, and suggested that the resulting transmembrane protein was potentially located on the surface of the plasma membrane of male gametes. A brief analysis of *PDI-Trans* is described within [37], where following a BarSeq Screen for asexual growth on an extensive library of non-clonal *P. berghei* KO parasites, it was posited that the gene is dispensable for the progression of blood-stage parasitemia. Our subsequent analysis of transcription levels by RT-PCR support this, demonstrating that *PDI-Trans* transcripts were present in wild-type asexual erythrocytic stages of gametocyte deficient strain 2.33, in addition to inactive (Gc-) and activated (Gc+) gametocytes, ookinetes and sporozoites (Figure 1A). To investigate the cellular localization of *PDI-Trans* across the parasitic lifecycle targeted-single homologous recombination was utilized to generate a transgenic *P. berghei* parasite expressing the endogenous *PDI-Trans* protein with a C-terminal EGFP fusion tag. Successful integration following drug selection was confirmed by PCR (Figure 1B). The presence of the EGFP tag caused no observable impact on blood or sexual stages, and did not impact transmission through *An. stephensi* mosquitoes. Immunofluorescence microscopy on non-permeablized parasites confirmed *PDI-Trans*-GFP expression on the surface of activated male gametes and ookinetes (Figure 1C). Live microscopy of mixed blood stages and fixed immunofluorescence of sporozoites demonstrated that *PDI-Trans*-GFP is expressed across the entire parasitic lifecycle (Figure S1).

**Figure 1.**
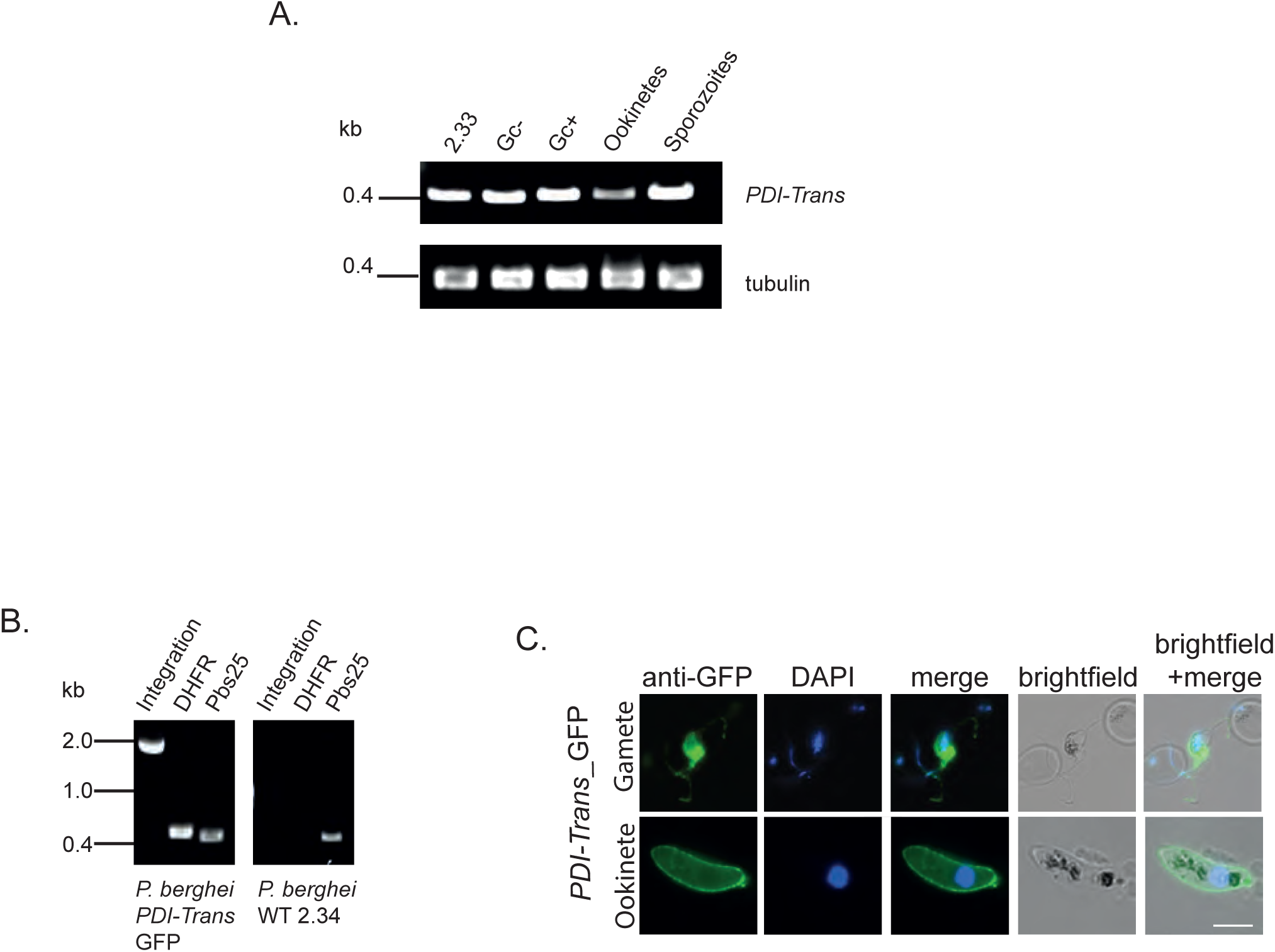
Constitutive expression of *Plasmodium berghei PDI-Trans*, and localization on the surface of gametocytes and ookinetes. ***A).*** RT-PCR analysis of *PDI-Trans* in asexual blood stages using the non-gametocyte producing strain 2.33; non-activated (Gc-) and activated (Gc+) gametocytes; purified *in vitro* ookinetes and day 21 salivary gland dissected sporozoites. The analysis was complemented with alpha-tubulin loading controls ***B).*** PCR confirmation of integration of *egfp* into the *PDI-Trans* locus. Oligonucleotides 35 and 14 were used to detect integration. Oligonucleotide 91 and 92 were used to detect DHFR presence, *pbs25* oligonucleotides were used as positive controls. *P. berghei* WT 2.34 gDNA was used as a negative control for integration. ***C).*** IFA of fixed, non-permeablised *PDI-Trans-GFP* parasites probed with anti-GFP; exflagellating male gametocytes (top) and ookinetes (bottom). Each panel shows an overlay of GFP fluorescence (green) and DNA labelled with DAPI (blue). White scale bar = 5 μm.

### PDI-Trans is essential for parasite transmission, is male specific and demonstrates classical PDI activity

To investigate the function of *PDI-Trans* targeted gene disruption was used to replace the entire *PDI-Trans* coding sequence. This was performed by double homologous recombination as described in [38.39], with constructs designed and manufactured by PlasmoGem (Sanger Institute, UK). Following dilution cloning of drug-resistant parasites, genotyping by PCR of two independently produced clones (Figure 2A) indicates that the replacement construct had integrated at the targeted site, disrupting the endogenous locus. Consistent with previous predictions [37], examination of mice infected with *ΔPDI-Trans* clones showed that the parasites underwent normal asexual development in erythrocytes (Figure S2). Rates of gametocytogenesis and sex ratio were unaffected, and gametocytes were able to emerge from their host cells and differentiate into gametes when exposed to standard gamete activation conditions (i.e. drop in pH or temperature, presence of xanthaurenic acid). To examine for a specific role during fertilization we specifically examined *in vitro* ookinete formation in blood collected from mice infected with *ΔPDI-Trans* parasites. Blood cultures from mice infected with *ΔPDI-Trans* parasites failed to produce ookinetes, a finding confirmed by triplicate experiments on two independent *ΔPDI-Trans* clones (Figure 2B). To further explore this phenotype *in vivo, An. stephensi* mosquitoes were fed on mice infected with *ΔPDI-Trans* parasites in triplicate, and 12 days later microscopy was used to examine the presence of oocysts (Figure S3). Triplicate experiments of each clone showed a mean reduction of 94.38% inhibition in intensity and 63.68% inhibition in prevalence with in *ΔPDI-Trans* clone 1, and a 96.43%/65.62% inhibition in intensity/prevalence with *ΔPDI-Trans* clone 2 when compared to wild type *P. berghei* (Table 1). These results suggest that *Trans-PDI* plays a key role in the successful transmission of *Plasmodium.*

**Figure 2.**
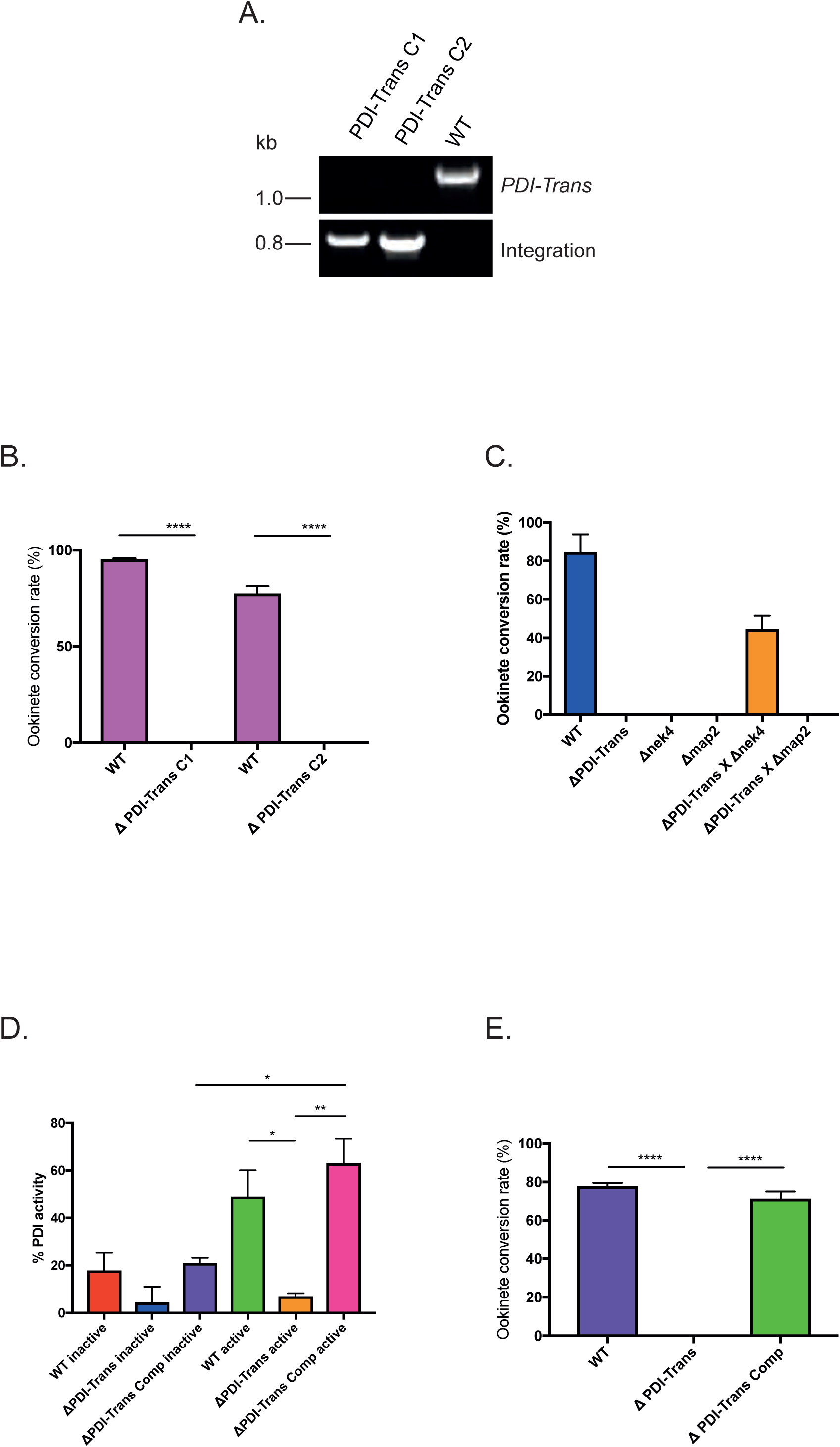
Deletion of *PDI-Trans* strongly inhibits transmission and is male specific. ***A).*** Diagnostic PCR with genomic DNA templates and primers 69 and 70 to test for the presence of *PDI-Trans*, and primers 72 and 9 to detect a unique 930bp product across the integrations site. ***B).*** The bar chart shows ookinete conversion rates for wild type and both *ΔPDI-Trans* clones. Conversion rate is expressed as a percentage of P28-positive parasites that had progressed to the ookinete stage (error bar indicates SEM; *n*=3). Asterisks indicate P value < 0.05 Paired t test ***C).*** *In vitro* ookinete conversion analysis demonstrates that *PDI-Trans* mutant shows production cross-fertilisation with the *Δnek4* sterility mutant, which produces functional males only, and not with *Δmap2* mutant, which produces functional females only. (error bar indicates SEM; *n*=3). Asterisks indicate P value < 0.05 Paired t test. ***D).*** PDI activity of purified active and inactive gametocytes from wild type, *ΔPDI-Trans* and *ΔPDI-Trans* Comp parasite lines. PDI activity is expressed as a percent relative to the positive control (human recombinant PDI). Experiments were performed in triplicate (error bar indicates SEM; *n*=3). Asterisks indicate P value < 0.05 Paired t test. ***E).*** Bar chart of ookinete conversion rates for wild type *ΔPDI-Trans* and *ΔPDI-Trans* Comp parasite lines. Conversion rate is expressed as a percentage of P28-positive parasites that had progressed to the ookinete stage (error bar indicates SEM; *n*=3). Asterisks indicate P value < 0.05 Paired t test

**Table 1.**
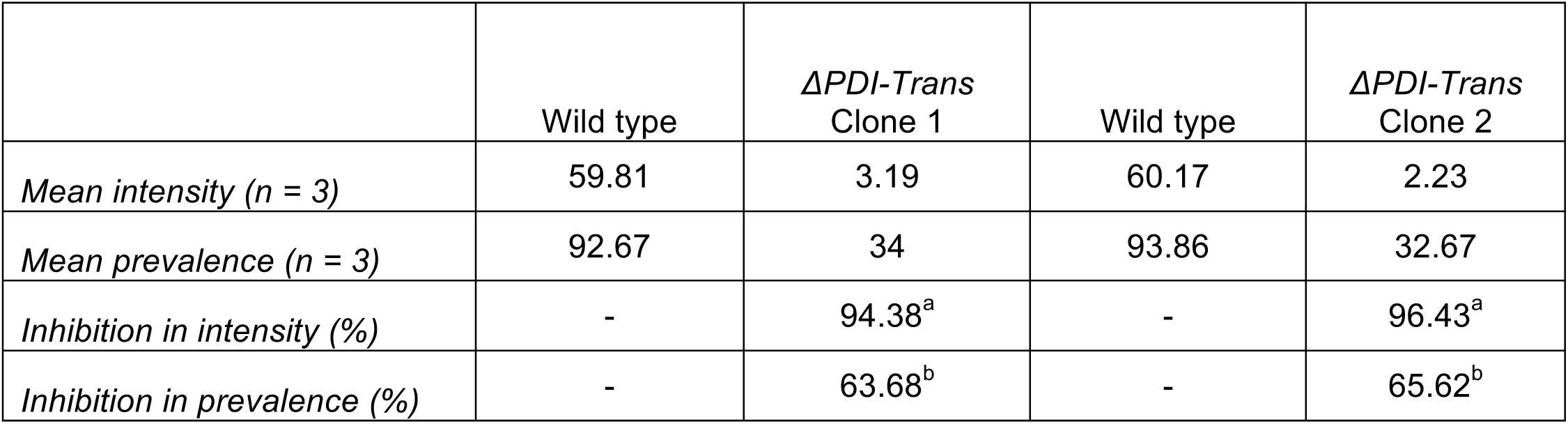
Mean *in vivo* evaluation of deleting *PDI-Trans* on transmission by direct feeding. The mean (from three replicates) inhibition in intensity (mean number of oocysts per midgut) and prevalence of two independent *PDI-Trans* knockout clones were calculated with respect to wild type controls. ^a^ P < 0.05, Mann-Whitney *U* test ^b^ P < 0.05, Fisher’s exact test.

|  | Wild type | *ΔPDI-Trans*<br>Clone 1 | Wild type | *ΔPDI-Trans*<br>Clone 2 |
| --- | --- | --- | --- | --- |
| *Mean intensity (n = 3)* | 59.81 | 3.19 | 60.17 | 2.23 |
| *Mean prevalence (n = 3)* | 92.67 | 34 | 93.86 | 32.67 |
| *Inhibition in intensity (%)* | - | 94.38<sup>a</sup> | - | 96.43<sup>a</sup> |
| *Inhibition in prevalence (%)* | - | 63.68<sup>b</sup> | - | 65.62<sup>b</sup> |

Cross-fertilization experiments, with known gender-specific sexual mutants, such as the male-deficient *map2* or the female-defective *nek4* mutant [40,41], make it possible to detect gender-specific sterility phenotypes in *P. berghei*. As shown in Figure 2C, neither *Δmap2* nor *Δnek4* strains produce ookinetes when cultured in isolation, but when cultures containing both KO lines were mixed, *Δnek4* male gametes are able to fertilize *map2* female gametes, restoring the capacity to form ookinetes (Figure 2C). Reduced conversion rates (compared with wild type parasites) are expected [3, 40, 41], due to the persistence of *Δnek4* female and *Δmap2* gametes which are unable to fertilize. In *ΔPDI-Trans /Δnek4* crosses the *PDI-Trans* female gametes were fertilized by *Δnek4* male gametes, but *Δmap2* females remained unable to differentiate into ookinetes in *ΔPDI-Trans/Δmap2* crosses (Figure 2C), indicating that *ΔPDI-Trans* males are sterile. Thus, these results demonstrate that during plasmodial fertilization, *PDI-Trans* is essential for microgamete (male) fertility.

PDI enzymes typically catalyze the rearrangement of disulphide bonds between cysteine residues within proteins. To determine whether *PDI-Trans* exhibits classical PDI activity we utilized a fluorescent PDI insulin-reduction assay to determine reductase activity. Recombinant human PDI was used as a positive control for PDI activity, and the well-characterized PDI inhibitor bacitracin was used as a negative control. Gametocytes from *ΔPDI-Trans, ΔPDI-Trans Comp* and wild type lines were purified on a density gradient and used within the assay in either an inactive, or activated form. PDI activity was expressed as a percentage relative to the positive control (Figure 2D). In wild type parasites, PDI activity is increased post-activation, implicating broad PDI activity throughout gamete activation/fertilization. The activated gametes of the *ΔPDI-Trans* line had significantly reduced PDI activity with respect to wild type gametes, specifically indicating that *PDI-Trans* exhibits true PDI reductase function during fertilization. PDI activity was significantly increased when *ΔPDI-Trans* was complemented with the endogenous gene *(ΔPDI-Trans Comp).* To investigate whether complementation restored not only PDI activity, but the ability of these parasites to successfully fertilize we performed ookinete conversion assays. Wild type parasites had a mean conversion rate of 77.98%. Ookinete conversion was not observed in the *ΔPDI-Trans* parasite line. Conversely, *ΔPDI-Trans Comp* parasites exhibited a mean ookinete conversion rate of 72.25% indicating that complementation of the *PDI-Trans* restored the ability to of male gametes to fertilize. Assays were performed in triplicate (Figure 2E).

### Malarial transmission is inhibited reversibly by the PDI inhibitor Bacitracin in P. berghei and P. falciparum

To further explore *PDI-Trans* activity, and to examine the ability of specific PDI inhibitors to block malarial transmission, we utilized the classical PDI inhibitor bacitracin [42]. The addition of bacitracin during fertilization *in vitro* resulted in a dose-dependent reduction of observable exflagellation centres (motile male and female gametes bound to each other), with a complete inhibition in formation of exflagellation centers at 3 mM (Figure 3A). This subsequently inhibited the ability of mature gametocytes to form ookinetes, with complete inhibition of ookinete conversion at 3 mM bacitracin (Figure 3B). In order to test whether bacitracin has a broad and non-specific toxic effect on parasites, potentially unrelated to *PDI-Trans* function, gametocytes were pre-incubated in bacitracin at a range of concentrations for 30 minutes, washed to remove the PDI inhibitor, then assayed for formation of exflagellation centers/ookinetes respectively (post-wash). Results show that parasites pre-incubated with bacitracin, then washed, resulted in no significant difference (Paired t test) in the number of exflagellation centres compared to the untreated control across all concentrations examined (Figure 3C-D).

**Figure 3.**
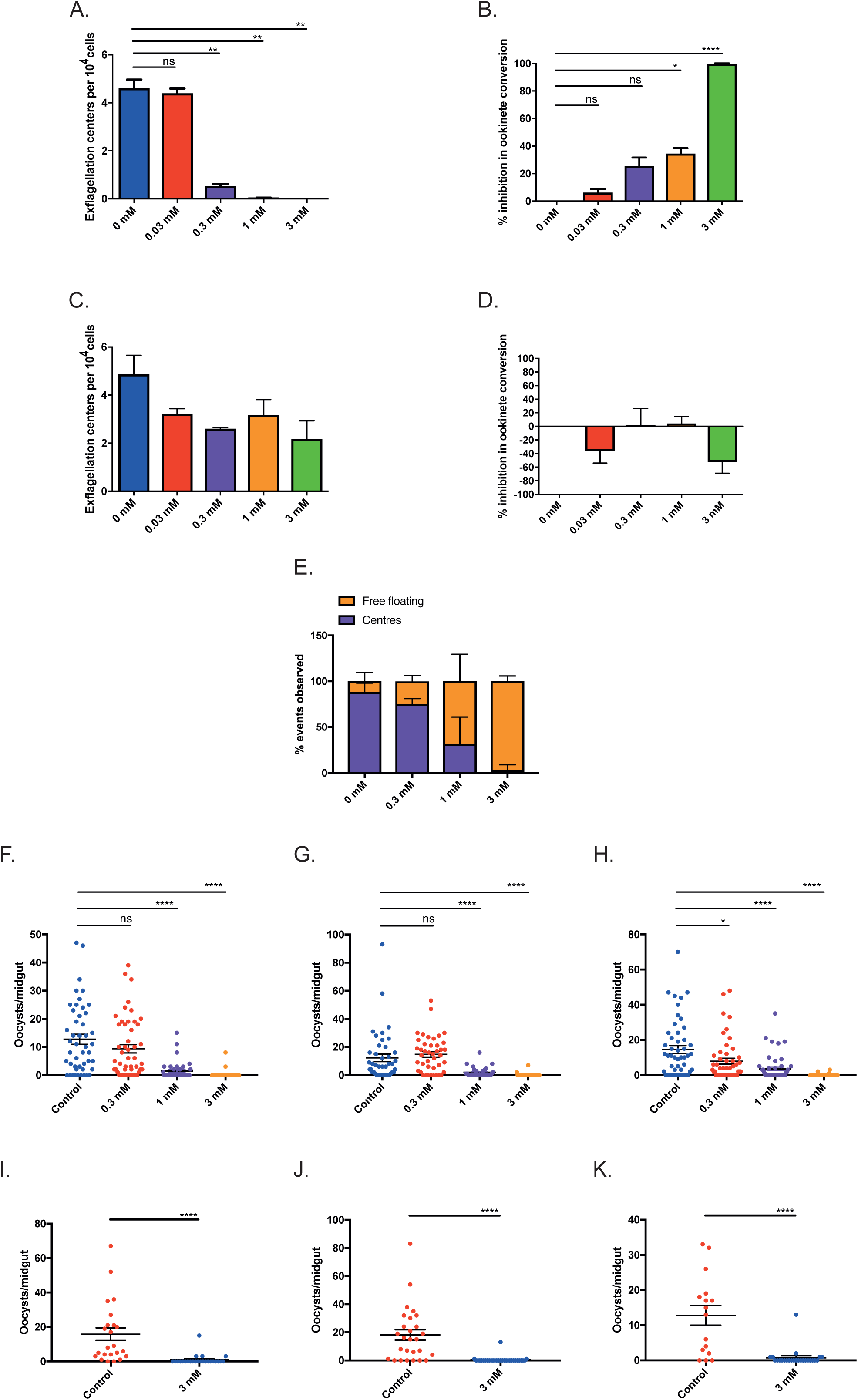
The specific PDI inhibitor bacitracin reversibly inhibits *Plasmodium berghei* fertilization *in vivo* and transmission *ex vivo*. ***A).*** Exflagellation centers in the presence of Bacitracin at 0, 0.03, 0.3, 1 mM. Asterisks indicate P value < 0.05 Paired t test. ***B).*** *In vitro* ookinete development assay supplemented with Bacitracin at 0, 0.03, 0.3, 1 and 3 mM. Results are shown as percent inhibition in ookinete conversion. Asterisks indicate P value < 0.05 Paired t test, ns indicate P value not significant. ***C).*** Exflagellation centers after 30 minutes in the presence of Bacitracin at 0, 0.03, 0.3, 1. P value < 0.05 Paired t test indicates P value not significant at any concentration. ***D).*** *In vitro* ookinete development assay supplemented with Bacitracin at 0, 0.03, 0.3, 1 and 3 mM for 30 min prior to removal of Bacitracin. Results are shown as percent inhibition in ookinete conversion. P value < 0.05 Paired t test indicates P value not significant at any concentration. ***E).*** Triplicate counts of free floating male gametes and male gametes in exflagellation centers. Represented as a percentage of total events observed in the presence of 0, 0.3, 1 and 3 mM Bacitracin. ***F-H).*** Triplicate *P. berghei* standard membrane feeding assays with Bacitracin compared to control at concentrations of 0.3, 1 and 3 mM. Individual data points represent the number of oocysts found in individual mosquitoes 12-days post feed. Horizontal bars indicate mean intensity of infection, whilst error bars indicate S.E.M within individual samples. Asterisks indicate P value <0.05 Mann-Whitney *U* test, ns indicate P value not significant. ***I-J).*** Triplicate *P. falciparum* standard membrane feeding assays with Bacitracin compared to control at a concentration of 3 mM. Individual data points represent the number of oocysts found in individual mosquitoes 8-days post feed. Horizontal bars indicate mean intensity of infection, whilst error bars indicate S.E.M within individual samples. Asterisks indicate P value <0.05 Mann-Whitney *U* test.

In an attempt to further examine the activity of *PDI-Trans* and the mechanism of PDI-inhibitor based blockade of transmission, following bacitracin treatment we examined the number of visible free floating male gametes present, compared with the number of exflagellation centers present (defined as three or more male gametes adhered to a female gamete). As bacitracin concentrations increased. the number of visible exflagellation centres decreased, conversely, the number of free floating observed microgametes was unaffected (Figure 3E), suggesting that PDI activity is essential for the association of male and female gamete to form exflagellation centers, but not for the ability of gametogenesis/activation to form microgametes.

To test the ability of bacitracin to block the transmission of *P. berghei ex vivo* we performed standard membrane feeding assays (SMFA) in triplicate with 0.3, 1 and 3 mM doses of bacitracin (Figure 3 F-H). Bacitracin inhibited transmission at all concentrations in a dose-dependent manner with a maximal inhibition in oocyst intensity of 98.21% and infection prevalence of 92.48% with 3 mM bacitracin (Table 2).

**Table 2.** Overall evaluation of transmission blocking effect of PDI inhibitor bacitracin in *P. berghei* by SMFA. Mean (from three replicates) reductions in intensity (mean number of oocysts per midgut) and prevalence with bacitracin at 0.3, 1 and 3 mM were calculated with respect to control feeds. ^a^ P < 0.05, Mann-Whitney *U* test ^b^ P < 0.05, Fisher’s exact test.

|  | Control | 0.3 mM | 1 mM | 3 mM |
| --- | --- | --- | --- | --- |
| *Mean intensity (n = 3)* | 13.17 | 10.68 | 2.32 | 0.23 |
| *Mean prevalence (n = 3)* | 77.37 | 72.32 | 38.29 | 5.81 |
| *Inhibition in intensity (%)* | - | 17.20<sup>a</sup> | 82.77<sup>a</sup> | 98.21<sup>a</sup> |
| *Inhibition in prevalence (%)* | - | 6.72<sup>b</sup> | 50.23<sup>b</sup> | 92.48<sup>b</sup> |

To investigate if PDI function is implicated in fertilization in additional *Plasmodium* species, and to extend our observations in *P. berghei* to human malaria parasites, we performed SMFAs with *P. falciparum* gametocyte cultures in the presence of bacitracin. Given the results observed with *P. berghei* we chose to perform the *P. falciparum* feeds at the highest concentration of bacitracin (3 mM) in triplicate to detect maximal effect (Figure 3 I-K). The addition of bacitracin significantly inhibited transmission, with a mean inhibition in oocyst intensity of 95.05% and an inhibition of oocyst prevalence of 81.71% observed (Table 3).

**Table 3.** Mean *ex vivo* evaluation of transmission blocking effect of PDI inhibitor bacitracin in *P. falciparum*. The mean (from three replicates) changes in intensity (mean number of oocysts per midgut) and prevalence with Bacitracin at 3 mM were calculated with respect to control feeds. ^a^ P < 0.05, Mann-Whitney *U* test ^b^ P < 0.05, Fisher’s exact test.

|  | Control<br>Feed 1 | Bacitracin<br>Feed 1 | Control<br>Feed 2 | Bacitracin<br>Feed 2 | Control<br>Feed 3 | Bacitracin<br>Feed 3 |
| --- | --- | --- | --- | --- | --- | --- |
| *Mean intensity* | 15.83 | 0.96 | 18.15 | 0.52 | 12.81 | 0.76 |
| *Mean prevalence* | 91.30 | 16 | 78.57 | 10 | 81.25 | 20 |
| *Inhibition in intensity (%)* | - | 93.93 | - | 97.14 | - | 94.07 |
| *Inhibition in prevalence (%)* | - | 82.48 | - | 87.27 | - | 75.38 |

### PDI-Trans can be targeted specifically with antibodies to block transmission in vitro and ex vivo

Considering the surface localization of *PDI-Trans* on the surface of the microgamete and ookinete, the ability of anti-*PDI-Trans* antibodies to initiate a specific anti-parasitic transmission blocking response was additionally examined. A polyclonal peptide antibody raised against residues bioinformatically predicted to be within the extracellular domain of *PDI-Trans* (amino acids 30 - 43). Antibodies were raised in rabbits, IgG purified, and examined for their ability to inhibit transmission *in vitro* and *ex vivo*. Anti-*PDI-Trans* IgG recognized both non-permablized gametocyte and ookinetes by IFA (Figure 4A). Staining was absent in secondary only controls, indicating the ability of these antibodies to specifically recognize natively folded *PDI-Trans* on the gamete and ookinete surface (Figure 4B).

**Figure 4.**
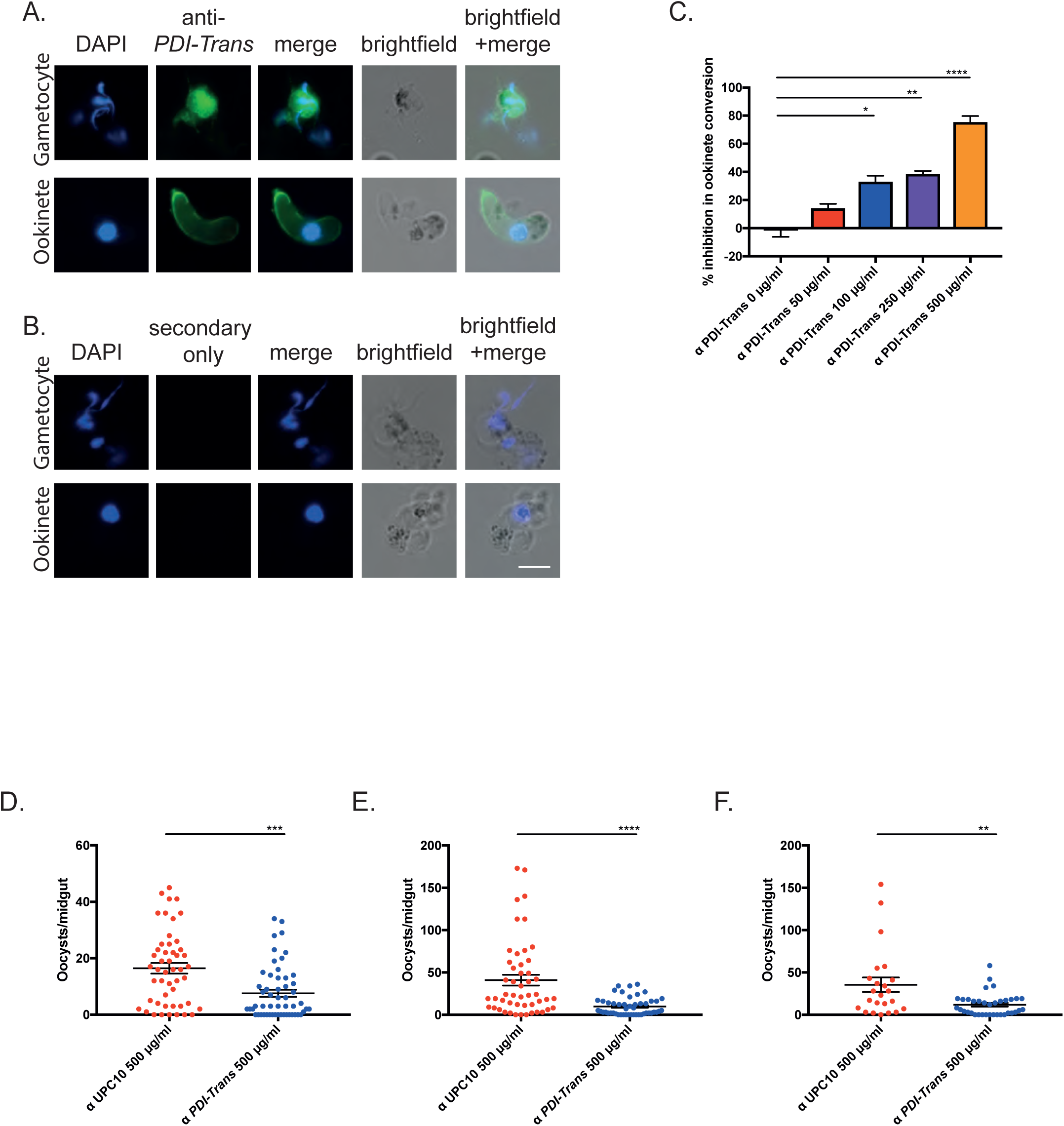
Anti-*PDI-Trans* antibodies inhibit fertilization and transmission in *Plasmodium berghei*. IFA of WT *P. berghei* ANKA male gametes and ookinetes with ***A).*** anti *PDI-Trans* and ***B).*** Secondary-only control antibodies (green) DAPI (blue). IFA of male gametes and ookinetes with anti *PDI-Trans* shows broad surface staining. White scale bars = 5 μm **C).** Inhibition in ookinete conversion in *in vitro* ookinete development assay with anti *PDI-Trans* antibody compared to negative control antibody UPC10 at concentrations of 0, 50, 100, 250 and 500 µg/ml. Asterisks indicate P value < 0.05 Paired t test, ns indicate P value not significant. ***D-F).*** Triplicate standard membrane feeding assays with anti *PDI-Trans* antibody compared with negative control antibody UPC10 at a concentration of 500 µg/ml. Individual data points represent the number of oocysts found in individual mosquitoes 12-days post feed. Horizontal bars indicate mean intensity of infection, whilst error bars indicate S.E.M within individual samples. Asterisks indicate P value <0.05 Mann-Whitney *U* test, ns indicate P value not significant.

To examine the ability of *PDI-Trans* to act a transmission blocking antigen, we performed *in vitro* ookinete conversion assays in the presence of anti-*PDI-Trans*. Anti-*PDI-Trans* inhibited ookinete conversion in a dose-dependent manner, further suggesting specificity. At antibody concentrations of 50, 100, 250 and 500 μg/ml, ookinete formation was inhibited by 14.3%, 33.2%, 38.7% and 75.4% respectively. In contrast, as previously demonstrated [4], the presence of UPC10 (negative control) had no effect on ookinete conversion (Figure 4C).

The transmission blocking activity of anti-*PDI-Trans* antibodies were additionally assessed by triplicate SMFA (Figure 4D-F). Given the *in vitro* results observed previously (Figure 4D), we assessed the *in vivo* transmission blocking ability of these antibodies only at the highest concentration where an effect in the *in vitro* ookinete assay was demonstrated. Anti-*PDI-Trans* antibodies significantly inhibited *P. berghei* transmission in all experiments. At a concentration of 500 μg/ml anti-*PDI-Trans* antibodies inhibited oocyst intensity by a mean of 66.22% and reduced prevalence of infection by 33.16% (Table 4).

**Table 4.** Mean *ex vivo* evaluation of transmission blocking effect of anti-*PDI-Trans* antibodies. The mean (from three replicates) change in intensity (mean number of oocysts per midgut) and prevalence with anti *PDI-Trans* antibody at 500 µg/ml were calculated with respect to appropriate negative control antibody UPC10 at the same concentration. ^a^ P < 0.05, Mann-Whitney *U* test ^b^ P < 0.05, Fisher’s exact test

| | $\alpha$ UPC10<br>500 μg/ml | $\alpha$ PDI-Trans<br>500 μg/ml |
| --- | --- | --- |
| *Mean intensity (n = 3)* | 31.37 | 9.82 |
| *Mean prevalence (n = 3)* | 93.28 | 62.18 |
| *Inhibition in intensity (%)* | - | 66.22<sup>a</sup> |
| *Inhibition in prevalence (%)* | - | 33.16<sup>b</sup> |

## Discussion

Fertilization is a key process in the *Plasmodium* lifecycle, encompassing the active fusion of activated male (micro) and female (macro) gametes to form a zygote within the mosquito bloodmeal. The resulting diploid zygote develops into a motile ookinete, establishing infection in the mosquito midgut and further progression throughout the Anopheline host. Despite its essential nature to the success of the parasitic lifecycle, the cellular and molecular mechanisms that underlie gamete fertilization of male and female gametes remain largely opaque in *Plasmodium* (and for the majority of Apicomplexa). Gamete interaction is a two-phase process; in the first phase, cell adhesion molecules displayed on the surfaces of the male and female gametes are responsible for gamete-gamete recognition. This initial recognition/adhesion step initiates a signal transduction cascade that activates the sperm and exposes new, fusogenic regions of the sperm plasma membrane. In *Plasmodium*, only three proteins have been discovered that have a demonstrable role in the mutual recognition of gametes; the 6-Cys family members, P48/45, P47 and P230 [5]. The specific mechanism of action of these proteins are currently unknown. In the absence of many of these surface proteins, there is a significant reduction in the formation of zygotes, however, low levels of fertilization still occur, particularly *in vitro* [3,6,7], indicating that gametes potentially use alternative, currently unknown, molecules to recognize each other. In the second phase of fertilization, the plasma membranes of the two gametes come into intimate contact and then fuse, bringing about cytoplasmic continuity. The conserved class II fusion transmembrane protein HAP2 is essential for gamete fusion during fertilization, and initiates merger of lipid bilayers post gamete adhesion. Following a (currently uncharacterized) trigger, it is hypothesized that a short-conserved region within the *Plasmodium* HAP2 ectodomain becomes exposed on the microgamete membrane surface, leading to the alignment of protein subunits parallel to each other, favoring trimerization [4,43,44]. Polar residues on the fusion loop are subsequently inserted into the target (macrogamete) membrane, followed by a conformational change in HAP2 domain III which distorts the target membrane, leading to hemifusion and then fusion/cytoplasmic continuity. Further related upstream and downstream effector molecules that specifically mediate the process of fertilization are at present unclear.

Here, we demonstrate that protein disulphide isomerase function, specifically encoded by a single plasmodial gene (*PDI-Trans*/PBANKA_0820300) is essential for malarial transmission. We demonstrate that *PDI-Trans* is constitutively expressed throughout the parasitic lifecycle, in both the blood and mosquito stages, but is only essential in the male gamete, where it is surface expressed. Absence of *PDI-Trans* only confers a detectable effect post-gamete activation, prior to gamete adhesion, and is null for male fertility, and consequently, zygote/ookinete formation, and transmission to the mosquito host. Complementation of the disrupted locus restores fertility. Furthermore, we conclusively demonstrate specific reductase activity, indicative of “classical” PDI activity, Inhibition of *PDI-Trans* using the widely available (topical) antibiotic and PDI inhibitor, bacitracin, reversibly blocks plasmodial transmission *in vivo* and *ex vivo*. Specifically, the process of gamete activation remains unaffected by bacitracin, whereas the ability of treated gametes to adhere to other cells (i.e. form exflagellation centers) appears to be compromised. Bacitracin-derived transmission blockade is observed in both *P. berghei* and *P. falciparum*. Finally, we show that antibodies specifically raised against the extracellular region of *PDI-Trans* can recognize the surface of the sexual stages of the parasite by immunofluorescence, and can initiate transmission-blocking activity both *in vitro* and *ex vivo*.

In all living cells, the appropriate formation and cleavage of disulphide bonds between cysteine residues in secreted and membrane-anchored proteins is essential for native conformation, and therefore, function. PDIs are traditionally known to be versatile enzymes with key roles in the mediation of disulfide bond formation, isomeration and reduction in the endoplasmic reticulum [32]. PDI function is also associated with varied chaperone activity [31]. Little is known regarding the expression and function of PDI-like proteins in *Plasmodium*. A previous study [33] has bioinformatically identified nine PDI-like molecules across five species of malaria parasites (four in *falciparum*, one in *vivax, berghei, knowlesi* and *yoelii*), indicated by the presence of classical thioredoxin domains. A more detailed analysis of one of these PDI candidates in *P. falciparum*; PfPDI-8 (PF3D7_0827900), demonstrated expression within the endoplasmic reticulum of asexual blood schizonts, gametocytes and sporozoites, with biochemical analysis indicating a function in the disulfide-dependent conformational folding of a recombinant form of the erythrocyte-binding protein (and putative bloodstage vaccine target) EBA-175. As further evidence of the chaperone function of PDI enzymes, studies utilizing the overexpression of PfPDI-8 resulted in the enhanced expression and folding of the transmission-locking vaccine candidate, Pf25, in *a Picha pastoris* expression system [59]. Broadly, PDI function is bioinformatically predicted to be conferred by multiple ORF throughout the parasitic genome. Expression, localization and function of these proteins are still largely undefined. Future study to further dissect the function of PDIs (and associated mechanisms of protein folding) in *Plasmodium* may be advantageous.

Although classically considered to be key mediators of protein folding in the endoplasmic reticulum, key evidence showing localization and function of PDIs in other cell compartments does exist. In some organisms, PDIs have been demonstrated to on occasion escape the ER, and exhibit cytoplasmic and cell surface localization, where their predominant function appears to be the reduction of disulphide bonds [32,45]. In terms of fertilization, PDI activity on the sperm head has previously proved to be essential for sperm-egg cell fusion in multiple vertebrates [31,46-48], and implicated in male fertility in mice [49.50]. PDI function has previously been implicated in the progression of multiple infectious diseases, with a specific role in mediating pathogen entry. In viruses, overexpression of PDI enhances the fusion of vial membranes, leading to increased internalization of HIV-1 [51]. Cell surface PDI has been shown to facilitate the infection of HeLa cells by mouse polyoma virus [52], and in endothelial cells a surface localized (lipid-raft associated) PDI reduces β1 and β2 integrins, allowing for the entry of dengue virus [53,54]. Its function is also essential for release of cholera toxin active chain A from the ER to the cytosol of the infected cell [54,55]. In protozoan pathogens, previous experimentation has demonstrated that increased levels of PDI increase the phagocytosis of the *L. chagasi* promastigote (but not the amastigote) [56]. It has previously been hypothesized that *T. gondii* and *L. donovani* PDI could be putative targets for vaccine development [57.58]. The specific function of *PDI-Trans* within the parasite is still unknown, however, it is clear that the process of successful fertilization in *Plasmodium* requires the presence and function of a range of proteins with conserved disulphide bonds between cysteine residues on the gamete surface. The 6-Cys family members P48/45, P47, P230 are all definitively evidenced to mediate gamete adhesion, whereas HAP2 requires the correct formation of multiple crucial disulphide bridges to enable membrane fusion. It cannot be discounted that *PDI-Trans* may in some way catalyse disulphide bond rearrangement in one of these transmission-essential proteins, exposing key residues critical for fertility-based function. Given the ability of the *PDI-Trans* knockout described here to undergo normal levels of gametogenesis, function post-activation, but pre-gamete fusion seems likely.

The results described here clearly indicate that *PDI-Trans* is a potential target for anti-malarial compounds or vaccines to successfully inhibit malarial transmission. The generation of novel TBIs to reduce disease burden is a key component of the current anti-malarial strategy, and it is widely accepted that to achieve eradication, it will be necessary to use interventions that inhibit the transmission of parasites from humans to mosquitoes [2]. We demonstrate that bacitracin reversibly inhibits malarial transmission with high efficacy, and additionally, that antibodies targeting *PDI-Trans* on the surface of the sexual stages of the parasite mediate significant transmission blocking immunity. It should be noted that bacitracin is already an FDA approved compound, traditionally used clinically against gram-positive compounds. This example illustrates the potential value of re-purposing drugs with observed efficacy against non-malarial species. Previous studies examining the anti-malarial efficacy of bacitracin only examined impact on asexual growth, where no effect was demonstrated [64]. The data here provides further evidence that potent anti-malarial transmission blocking efficacy can be achieved by targeting the male (micro) gamete. Previously described compounds effective against the process of fertilization (methylene blue and atovaquone) are effective in blocking transmission, as are antibodies against multiple male gamete-surface proteins [21-27]. To target the sexual stages of the malaria parasite further, a deeper understanding of transmission and specifically, the mechanism of fertilization within *Plasmodium* is advantageous, and offers the potential for the development of new, effective interventions. More broadly, PDI-like proteins are expressed across multiple taxa, and species, including in a wide range of organisms of veterinary and clinical importance [31,46]. Given the ability of both anti*-PDI-Trans* compounds and antibodies to block malarial transmission described here, and the proven role of PDI function in the regulation of infection across multiple species, it is not unreasonable to suggest that further studies may want to examine the possibility of targeting PDI proteins/functions using specifically designed novel anti-malarial drugs or vaccines.

## Experimental procedures

### General parasite maintenance

General parasite maintenance was carried out as described in [60]. Briefly, *P. berghei* parasites were maintained in 6–8-week-old female Tuck Ordinary (TO) mice (Harlan) by serial mechanical passage (up to a maximum of eight passages). If required, hyper-reticulosis was induced three days before infection by treating mice intraperitoneally (*i.p*) with 200 μl phenylhydrazinium chloride (PH; 6 mg/ml in PBS; ProLabo UK). Mice were infected *i.p.* and infections were monitored using Giemsa-stained tail blood smears as described previously [70].

### Generation and analysis of transgenic parasite lines

#### PDI-Trans-GFP

To examine the expression and localization of *PDI-Trans*, the *PDI-Trans*-GFP transgenic line was created, introducing a C-terminal GFP tag to the native by single homologous recombination. The targeting construct *pPDI-Trans-GFP* was constructed using the backbone of the EGFP-tagging vector p277 [62]. The terminal 1527 bp of the *PDI* gene (PBANKA_082030) was synthesized (IDT) to remove internal *<u>ApaI</u>* site and introduce unique *AvrII* site within the gene and flanking *KpnI* and *ApaI* sites to the amplicon. This block was cloned in frame into *ApaI*/*KpnI* sites of p277, resulting in *pPDI-Trans -GFP*. For transfection, this construct was linearized at a unique *AvrII* site within the *PDI* sequence.

Parasites were transfected using the Nucleofector device (Amaxa Biosystems) as described previously [62]. Integration of the DNA constructs into the chromosome was confirmed by PCR flanking a region upstream from the 5′ integration site into the EGFP sequence (oligo 35; 5′-GCATGTGCGATTGTATTGGG-3; oligo 14; 5′-ACGCTGAACTTGTGGCCG-3′) and the presence of the DHFR selection cassette (oligo 91 5’-TTCGCTAAACTGCATCGT −3’; oligo 92 5’-GTACTTAATGCCTTTCTCCT-3’). Oligos against the *Pbs25* gene (PBANKA_0051500) were used as positive control (oligosF1: 5′-CAACTTAGCATAAATATAATAATGCGAAAGTTACCGTGG-3′;F25′-CCATCTTTACAATCACATTTATAAATTCCATC-3′). GFP expression in transfected, drug resistant parasites were confirmed by fluorescence microscopy. Two independent clones were obtained from two independent transfections, demonstrating identical phenotypes and GFP expression.

#### ΔPDI-Trans

To examine the function of PBANKA_082030 the *ΔPDI-Trans* transgenic line was generated. The plasmid was designed and constructed by PlasmoGEM (PlasmoGem ID PbGEM-239637) using recombinase-mediated engineering followed by a Gateway® mediated exchange [39,63]. Prior to transfection the construct was digested by NotI to release the *P. berghei* insert from its vector backbone. Parasites were transfected using the Nucleofector device (Amaxa Biosystems) as described previously [62]. Integration of the DNA constructs into the chromosome was confirmed by PCR region flanking 5’ of the modified target locus and 3’ DHFR selection cassette (oligo 72; 5′-ACGTGCATGTGCGATTGTATTGGGT −3; oligo 9; 5′-CTTTGGTGACAGATACTAC −3′) and the absence of the wildtype locus (oligo 69 5’-ATGGGAAACTATACTTATATATATATTTTTTTCA −3’; oligo 70 5’-TTATAAATCAGAATTTTCTTCTCCTTC −3’). Two independent clones were obtained from two independent transfections, demonstrating statistically indistinguishable phenotypes.

#### ΔPDI-Trans-Comp

For the complementation construct the clonal knockout line was injected into mice and mice were treated with 5-Fluorocytosine (5FC) nucleoside analog (Sigma) drinking water, 1.5 mg/ml to recycle the *Hdhfr*-*yfcu* marker. The subsequent marker free line was subjected to dilution cloning to achieve a pure population of marker free parasites. Following this the full length endogenous PBANKA_082030 gene was transfected on top using the artificial chromosome library clone mapping to PBANKA_082030 from PlasmoGEM (clone ID PbAC02-74d11) as described previously (Figure S4) [37, 62].

### RT-PCR

*P. berghei* RNA was isolated from gametocyte deficient strain 2.33, activated or inactivated gametocytes, ookinetes and sporozoites from wild type *P. berghei* 2.34 strain using Trizol reagent (Invitrogen). cDNA synthesis was performed using Prime script kit from (Clonetech). PCR reactions were set up to amplify sections of *PDI-Trans* ORF (Forward 5’-ATGGGAAACTATACTTATATATATATTTTTTTCA-3’; and reverse 5’-CTACATATTTATCGACATCTCCAA-3’). The expected RT amplicon was 481 bp. The ubiquitously expressed α-tubulin gene PB300720.00.0 was amplified for each sample to ensure amplifiability of cDNA from respective RNA samples (Forward, 5′-CCAGATGGTCAAATGCCC-3′; Reverse, 5′-CTGTGGTGATGGCCATGAAC-3′). The expected products were 435 bp (cDNA). Thirty RT–PCR cycles were carried out with denaturation for 1 min at 94°C, annealing for 45 secs at 50°C, and extension for 1.5 min at 68°C, and products were visualized on a 0.8% agarose gel.

### Direct Feeding Assay (DFA)

Routine maintenance of *P. berghei* was carried out as described above. Prior to challenge, mice were PH treated, and 3 days later infected *i.p.* with 10^6^ *P. berghei* ANKA 2.34 or *ΔPDI-Trans* parasites. Three-days post-infection, animals were anesthetized, and >50 female *Anopheles stephensi* mosquitoes allowed to blood feed on each mouse. Twenty-four hours later, unfed mosquitoes were removed. Mosquitoes were maintained on 8% (w/v) fructose, 0.05% (w/v) p-aminobenzoic acid at 19-22 °C and 50-80% relative humidity. Day 14 post-feeding, mosquito midguts were dissected and oocyst intensity and prevalence observed by standard phase microscopy and recorded. Reduction in oocyst intensity and prevalence in knockout mice were calculated with respect to wild type controls.

### In Vitro Ookinete Conversion Assay (IVOA)

PH-treated mice were injected with 5×10^7^ parasites *i.p.* On day 3 or 4 of infection, parasitaemia was counted on a Giemsa-stained tail blood smear and exflagellation of male gametocytes was checked by addition of a drop of exflagellation medium to a drop of tail blood. Hosts observed to have exflagellating parasites were exsanguinated by cardiac puncture and each 20 μl of blood taken up in 450 μl ookinete medium. Individual cultures were then added to pre-prepared 24 well plates (Nunc) and incubated for 24h at 19°C. Cultures were harvested after 24h by centrifugation (500×g, 5min), washed once in 100 μl ookinete medium, and the pellet taken up in 50 μl ookinete medium containing Cy3-conjugated Pbs28 mAb clone 13.1 (1:500). Ookinetes and macrogametocytes were then immediately counted by fluorescence microscopy. Ookinete conversion rates were calculated as described previously [19]. In bacitracin experiments harvested parasites were added to ookinete medium containing a range of Bacitracin (Sigma Aldrich: #B0125) concentrations and either left in or washed and put into fresh medium 30 min after drug treatment. In antibody experiments harvested parasites were added to ookinete medium containing anti-*PDI-Trans* rabbit sera or anti-UPC10 (negative control). In each set of experiments results were collated from three separate experiments and inhibition expressed as the percentage reduction in ookinete conversion with respect to wild type parasites, samples with no bacitracin or the anti-UPC10 control.

### Crosses

At day 3 post infection of phenylhydrazine treated mice, infected with parasites with either *ΔPDI-Trans, Δnek4, Δmap2* and wt were harvested by heart puncture and mixed at a 1:1 ratio in ookinete medium. After 24 h, ookinete conversion assays were performed by incubating samples with 13.1 antibody (antibody against Pb28 conjugated with Cy3). The proportion of ookinetes to all 13.1-positive cells (unfertilised macrogametes and ookinetes) was established, counting fields at 60 × magnification. Experiments were performed in biological triplicate [40,41].

### PDI activity assay

PDI activity was measured in a microplate PDI inhibitor screening assay kit from Abcam (ab139480). Briefly, *ΔPDI-Trans ΔPDI-Trans* Comp and wild-type gametocytes were purified [72]. Both activated and non-activated gametocytes of each parasite line were used in the assay and the PDI-catalyzed reduction of insulin in the presence of Dithiothreitol resulting in the formation of insulin aggregates which bind avidly to the red-emitting fluorgenic PDI detection reagent were measured on Tecan, Infinite M200 Pro. The background media signal for each sample was subtracted and PDI activity was calculated as a percent relative to the positive control (human recombinant PDI). Experiments were performed in triplicate.

### Standard Membrane Feeding Assay (SMFA)

#### P. berghei

Female *An. stephensi* (SDA 500 strain) were starved for twenty-four hours and then fed on heparinized *P. berghei* infected blood using standard membrane feeding methods [60]. For each feed, 350 µl of *P. berghei* ANKA 2.34 infected blood containing asexual and sexual stages of the parasite was mixed with 150 µl of PBS containing either antibody to yield final concentration of 500 μg/ml or drug at 0.3, 1 and 3 mM. Mosquitoes were handled, maintained and analyzed as described above. Reductions in oocyst intensity and prevalence was calculated with respect to control feeds as described in [4].

#### P. falciparum

Mature gametocytes of *P. falciparum* (NF54) were produced *in vitro* as described previously [65] with slight modifications. Briefly, mature gametocyte cultures (0.5 to 2 % final gametocytaemia) were fed for 15-20 min at room temperature to *An. gambiae* mosquitoes through an artificial membrane kept at 37 °C. For each feed 300 µl of mature *P. falciparum* gametocytes were mixed with bacitracin at a concentration of 3 mM. Engorged mosquitoes were housed in pots at 26°C and 60–80% relative humidity. On days 7-9, midguts were dissected and the results analyzed as outlined in the above *P. berghei* section.

### Antibody production

Synthetic peptide to *PDI-Trans* (VSDDFAKKVNHLTHC) was produced, conjugated to KLH and used to raise polyclonal rabbit antisera (Genscript, USA). Resulting sera was IgG purified and validated by Genscript via ELISA.

### Microscopy

#### Immunofluorescence assay (IFA)

*PDI-Trans*-GFP parasites were assessed by IFA for the presence of GFP tag with anti-GFP, Roche at a dilution of 1:500. Signal was detected by Alexa Fluor 488-labelled goat, anti-mouse IgG (Molecular Probes) at 1:500. Rabbit antibodies to *PDI-Trans* were assessed by IFA on wild-type *P. berghei* ANKA 2.34 gametocytes and ookinetes at a dilution of 1:500. Signal was detected by Alexa Fluor 488-labelled goat, anti-rabbit IgG (Molecular Probes) at 1:500. Parasites were cultured and IFAs were performed as described previously [4]. Slides were visualized under x60 objective magnification using a fluorescence microscope (EVOSFL Cell Imaging System, Life Technologies).

#### Live imaging

*PDI-Trans*-GFP parasites were examined for GFP signal by live microscopy. Parasites were cultured and allowed to settle on glass slides before microscopy. Slides were visualized under X40 objective magnification using a fluorescence microscope (Leica DMR).

### Statistical Analysis

Statistical analysis was performed using Graphpad Prism. For DFA, SMFA and DMFA, significance was assessed using Mann–Whitney U (to examine differences in intensity) and Fisher’s exact probability tests (to examine differences in prevalence). Parametric ELISA tests were assessed using t-test. P values < 0.05 were considered statistically significant (^***^ = <0.0001, ^***^= 0.001, ^**^= 0.001-0.01, ^*^= 0.01-0.05).

### Ethical Statement

All procedures were performed in accordance with the UK Animals (Scientific Procedures) Act (PPL 70/8788) and approved by the Imperial College AWERB. The Office of Laboratory Animal Welfare Assurance for Imperial College covers all Public Health Service supported activities involving live vertebrates in the US (no. A5634-01).

## Acknowledgements

This work was funded by the MRC (New Investigator Research Grant; award number MR/N00227X/1) A.M.B thanks PATH-MVI for funding. Funders had no role in study design, data collection and interpretation, of the decision to submit the work for publication. We gratefully acknowledge Mark Tunnicliff for mosquito production and Tibebu Habtewold for advice regarding *P. falciparum* infections.

---

**Supplemental Figure S1.**
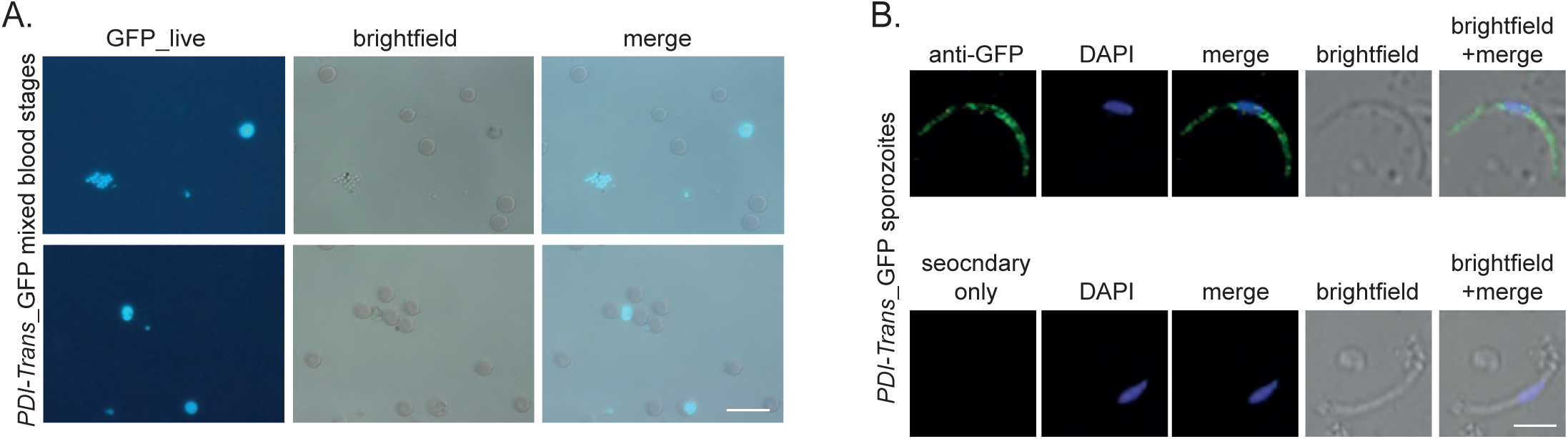
***A).*** Live GFP fluorescence of mixed blood stage *P. berghei PDI-Trans-GFP* parasites. Scale bar = 15 μm. ***B).*** IFA of fixed, non-permeablised *PDI-Trans-GFP* salivary gland sporozoites probed with either anti-GFP (top) or secondary only (bottom). Each panel shows an overlay of GFP fluorescence (green) and DNA labelled with DAPI (blue). Scale bar = 5 μm.

**Supplemental Figure S2.**
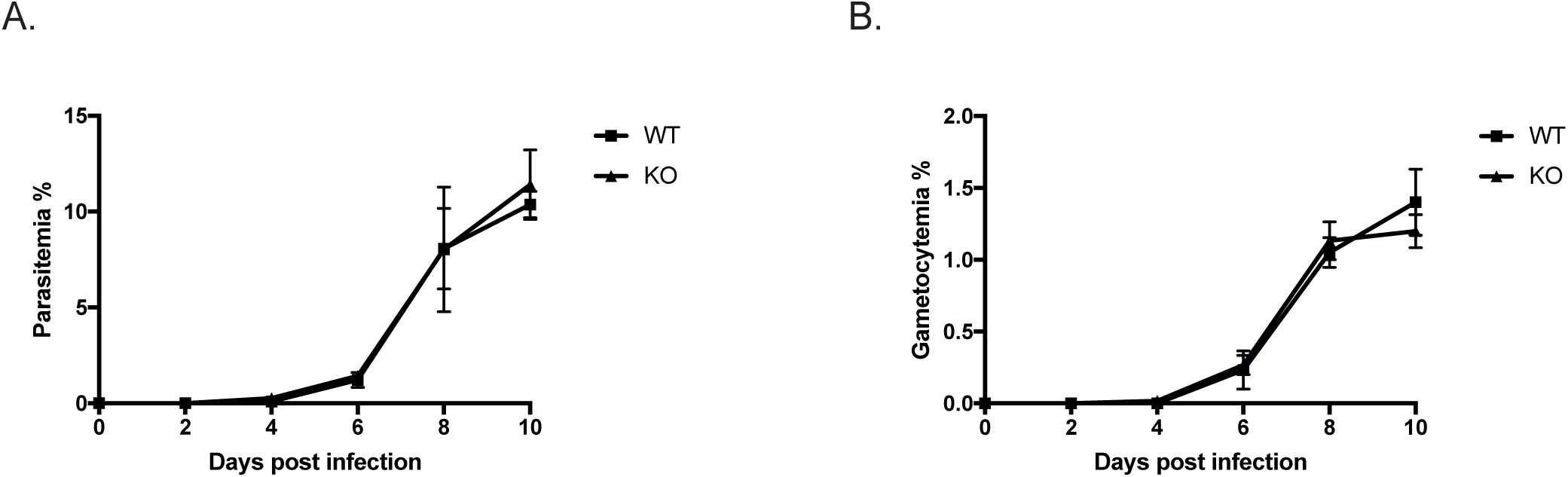
***A).*** Asexual growth and ***B).*** gametocyte production of WT and *ΔPDI-Trans* parasites strains. Three independent experiments are plotted.

**Supplemental Figure S3.**
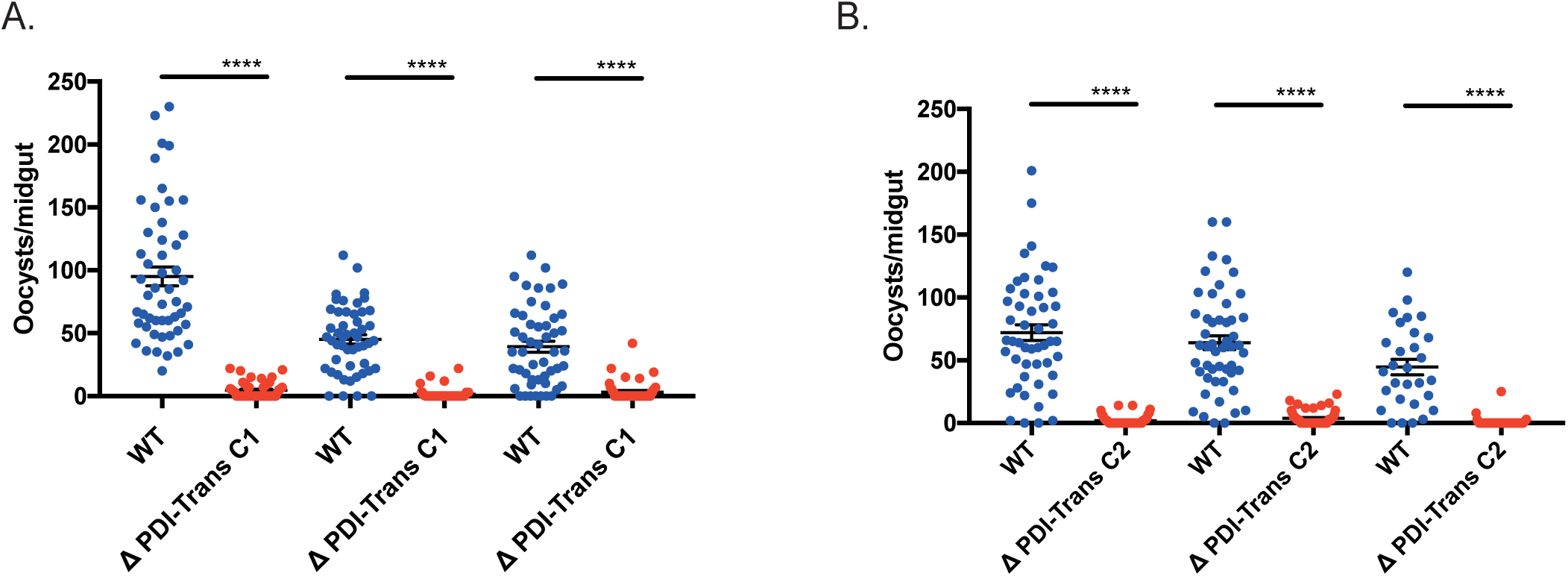
Mice infected with ***A).*** *ΔPDI-Trans* clone 1 (C1) or ***B).*** *ΔPDI-Trans* clone 2 (C2) *P. berghei* parasites and DFA performed to determine transmission blockade. Individual data points represent the number of oocysts found in individual mosquitoes 12 days post feeding. Horizontal bars indicate mean intensity of infection, while error bars indicate SEM within individual samples. Asterisks indicate P value < 0.05 Mann-Whitney U test.

**Supplemental Figure S4.**
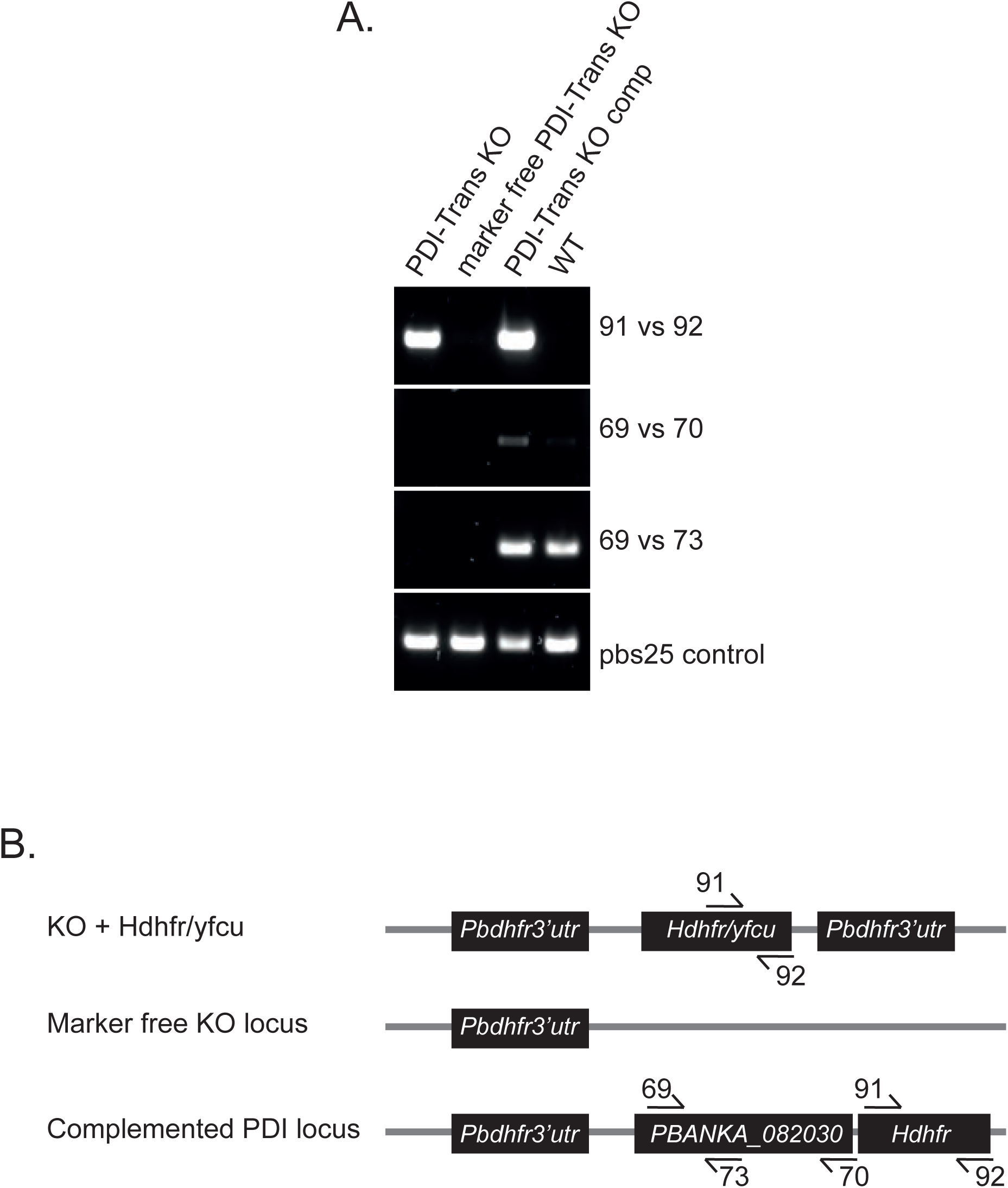
***A).*** Genotyping data for *ΔPDI-Trans, ΔPDI-Trans* marker-free and *ΔPDI-Trans* Comp lines. ***B).*** Schematic of for *ΔPDI-Trans, ΔPDI-Trans* marker-free and *ΔPDI-Trans* Comp lines and primer pairs used for genotyping.

